# Can malaria parasites manipulate the odour-mediated host preference of their mosquito vectors?

**DOI:** 10.1101/114272

**Authors:** Phuong L. Nguyen, Amélie Vantaux, Domonbabele FdS Hien, Kounbobr R. Dabiré, Bienvenue K. Yameogo, Louis-Clément Gouagna, Didier Fontenille, François Renaud, Frédéric Simard, Carlo Costantini, Fréderic Thomas, Anna Cohuet, Thierry Lefèvre

## Abstract

Malaria parasites can manipulate mosquito feeding behaviours such as motivation and avidity to feed on vertebrate hosts in ways that increase parasite transmission. However, in natural conditions, not all vertebrate blood-sources are suitable hosts for the parasite. Whether malaria parasites can manipulate mosquito host choice in ways that enhance parasite transmission toward suitable hosts and/or reduce mosquito attraction to unsuitable hosts (i.e. specific manipulation) is unknown. To address this question, we experimentally infected three species of mosquito vectors (*Anopheles coluzzii, Anopheles gambiae,* and *Anopheles arabiensis*) with wild isolates of the human malaria parasite *Plasmodium falciparum,* and examined the effects of immature (oocyst) and mature (sporozoite) infections on mosquito behavioural responses (activation rate and odour choice) to combinations of calf odour, human odour and outdoor air using a dual-port olfactometer. Regardless of parasite developmental stage and mosquito species, *P. falciparum* infection did not alter mosquito activation rate or their choice for human odours. The overall expression pattern of host choice of all three mosquito species was consistent with a high degree of anthropophily, with both infected and uninfected individuals showing higher attraction toward human odour over calf odour, human odour over outdoor air, and outdoor air over calf odour. Our results suggests that, in this system, the parasite may not be able to manipulate the early long-range behavioural steps involved in the mosquito host-feeding process, including initiation of host-seeking and host orientation. Future studies examining mosquito host-feeding behaviours at a shorter range (i.e. the “at-host” foraging activities) are required to test whether malaria parasites can modify their mosquito host choice to enhance transmission toward suitable hosts and/or reduce biting on unsuitable hosts.

## 1. Introduction

Host behavioural manipulation by parasites is a widespread transmission strategy (Moore, 2002; Lefèvre et al., 2009a; Poulin, 2010; Hughes et al. 2012). Trophically-transmitted parasites, for example, can alter the behaviour of their intermediate hosts in ways that increase predation rate by definitive hosts, hence favouring transmission (Lafferty and Morris, 1996; Lagrue et al., 2007; Lagrue et al., 2013). However, altering the behaviour of intermediate hosts can also increase predation rates by unsuitable hosts (Mouritsen and Poulin, 2003; Kaldonski et al., 2008; Seppälä et al., 2008). This higher probability of being killed by deadend predators can incur significant costs to manipulative parasites, especially when initial predation risk is high (Seppälä and Jokela, 2008). In response, some parasites have evolved specific manipulation, i.e. the ability to enhance transmission toward appropriate hosts and/or reduce predation by unsuitable hosts (Lagrue et al., 2007; Médoc and Beisel, 2009; Médoc et al., 2009; Seppälä et al., 2012; Lagrue et al., 2013; Jacquin et al., 2014).

In addition to its ecological and evolutionary relevance, host manipulation may also have profound implications for human health. Many manipulative parasites are responsible for devastating vector-borne diseases such as dengue fever, malaria, leishmaniasis, or sleeping sickness. Vector-borne parasites can indeed manipulate phenotypic traits of their vectors and hosts in ways that increase contacts between them, hence favouring parasite transmission (Hurd, 2003; Lacroix et al., 2005; Lefèvre and Thomas, 2008; Cator et al., 2012; Cornet et al., 2013a; De Moraes et al., 2014; Caljon et al., 2016). A frequently reported change induced by vector-borne parasites is alteration of vector motivation and avidity to feed. For example in malaria mosquitoes, individuals infected with *Plasmodium* sporozoites (the mosquito to human transmission stages) can display increased response to host odours (Rossignol et al., 1986; Cator et al., 2013), increased landing and biting activity (Rossignol et al., 1984, 1986;Wekesa et al. 1992; Anderson et al. 1999; Koella et al., 2002; Smallegange et al., 2013;), increased number of feeds (Koella et al., 1998) and increased blood volume intake (Koella and Packer, 1996; Koella et al., 1998; Koella et al. 2002). In contrast, mosquitoes infected with oocysts (the immature non-transmissible stage of the parasite), are less persistent and less likely to attempt to feed (Anderson et al., 1999; Koella et al., 2002; Cator et al., 2013). Since biting is risky (e.g., host defensive behaviours can kill the vector and its parasite), reduced feeding attempts seems beneficial to the parasite (Schwartz and Koella, 2001).

In natural conditions, these “stage-dependent” behavioural changes presumably increase the rate at which a mosquito will feed on a vertebrate blood-source, not all of which are suitable hosts for the parasite. The very few epidemiological models that have considered mosquito behavioural manipulation by malaria parasites have assumed that suitable hosts for parasite development were the only source of blood (Dobson, 1988; Cator et al., 2014). These models have ignored the possibility that malaria parasites may increase biting rate on vertebrates that do not act as suitable hosts. Similarly, no study has, to our knowledge, investigated whether malaria parasites can manipulate mosquito vertebrate choice in ways that enhance parasite transmission toward suitable hosts and/or reduce mosquito attraction to unsuitable hosts (i.e. specific manipulation).

Mosquito choice for vertebrate blood-source is an important key predictor for the transmission intensity of vector-borne diseases. This choice may be influenced by genetic and environmental factors such as the innate preference of the mosquito and the availability of the vertebrate species (Lyimo and Ferguson, 2009). While some malaria vectors can display propensity to feed on different vertebrate species (i.e. generalist or opportunistic feeding behaviour) (Takken and Verhulst, 2013), the parasites they transmit are often highly host-specific, infecting only one or a few vertebrate species (Perkins, 2014). Because of this strong host specificity, it is possible that vector-borne parasites acquired, during the course of evolution, the ability to target appropriate host and/or avoid unsuitable ones (Lefèvre et al., 2006). Accordingly, generalist mosquitoes, once infected, should develop a feeding preference for vertebrates that are suitable for parasite development. Studies exploring this possibility may yield important information about the diversity of transmission strategies used by malaria parasites and show that mosquitoes infected with transmissible parasite stages not only bite “more” but perhaps also “better”.

Theoretically, there are several ways through which malaria parasites could maximise transmission towards suitable vertebrate hosts. First, the parasite may induce in the mosquito vector a sensory bias for host traits (e.g. specific odours) that are correlated with optimal suitability for the parasite. Second, the parasite may induce alteration of mosquito microhabitat choice, in a way that spatially matches the microhabitat of the suitable host species. Finally, the parasite may induce changes in time activity in a way that temporally matches the resting time of the suitable host.

Here, we explored the first possibility using the natural association between *Plasmodium falciparum,* which causes the most severe form of human malaria, the mosquito species *Anopheles coluzzii, Anopheles gambiae* and *Anopheles arabiensis,* three major vectors of *P. falciparum* in Africa, and calves and human, two common mosquito vertebrate hosts. *An. coluzzii* and *An. gambiae* are considered anthropophilic (they are attracted to human stimuli) throughout its distribution, whereas *An. arabiensis* can display a weak host tropism (plastic/opportunistic) and can display either anthropophilic or zoophilic preference depending on the geographic area and the relative abundance of cattle and human (Costantini et al., 1999; Takken and Verhulst, 2013). *P. falciparum* displays an extreme form of specificity and can develop and reproduce in hominids only (predominantly in human and to a lesser extent in chimpanzee, bonobo, and gorilla) (Prugnolle et al., 2011; Rayner et al., 2011; Ngoubangoye et al., 2016), such that any mosquito bite on another vertebrate species, such as cattle, would be a dead-end for the parasite. We experimentally challenged local colonies of three mosquito species with sympatric field isolates of *P. falciparum* using direct membrane feeding assays in Burkina Faso, and examined the effects of immature (oocyst) and mature (sporozoite) infections on mosquito choice between human and calf odours using a dual-port olfactometer.

## 2. Material and methods

### 2.1. Mosquitoes

*Anopheles gambiae and An. coluzzii* mosquitoes originated from outbred colonies established in 2008 and repeatedly replenished with F1 from wild-caught mosquito females collected in Soumousso (*An. gambiae*) (11°01’00”N, 4°02’59’’W) and Kou Valley (*An. coluzzii*) (11°23’14”N, 4°24’42’’W), south-western Burkina Faso (West Africa) and identified by PCR-RFLP (Fanello et al., 2002; Santolamazza et al., 2008). Mosquitoes were maintained under standard insectary conditions (27 ± 2°C, 70 ± 10% relative humidity, 12:12 LD). Larvae were bred in the laboratory with *ad libitum* Tetramin^®^ and adult mosquitoes were provided with a solution of 5% glucose.

*Anopheles arabiensis* mosquitoes originated from wild caught larvae in Dioulassoba (11°10’42”N, 4°18’26’’W), a district of Bobo Dioulasso, where previous surveys ensured that *An. arabiensis* population was dominant (Dabiré et al., 2012, 2014). Collection of field larvae was conducted three times (twice in October 2014 and once in November 2014). Larvae were reared under the same standard insectary conditions as the mosquito colonies and F0 females were used for the experiments. Species identification of 35 individual from each wild caught batch (a total of 105 mosquitoes) was performed to confirm that *An. arabiensis* was the dominant species (Fanello et al., 2002). Samples from the 1^st^ and 2^nd^ batches in October contained 60% and 90.91% of *An. arabiensis* respectively; sample from the 3^rd^ batch in November contained 100% *An. arabiensis.*

### 2.2. Experimental infections

Experimental infections of mosquitoes were performed by membrane feeding of infectious blood (DMFA for Direct Membrane feeding Assay) as described previously (Ouédraogo et al., 2013; Vantaux et al., 2014; Vantaux et al., 2015; Roux et al., 2015; Hien et al., 2016). Briefly, three-to five-days old females were fed through membranes on *P. falciparum* gametocyte (the human to mosquito transmission stage)-infected blood from malaria patients in Burkina Faso. Mosquitoes were starved of glucose solution for 12 h (An. *gambiae* and *An. arabiensis*) or 24 h (*An. coluzzii*) prior to the infection. Gametocyte carriers were selected by examining thick blood smears from children aged between 5 and 11 from two villages in southwestern Burkina Faso (Dande and Soumousso, located 60km north and 40km southeast of Bobo Dioulasso, respectively) and blood drawing was carried out at laboratory. For *An. gambiae* and *An. arabiensis* mosquitoes, when gametocytemia was below 160 gametocytes/μl, blood serum was replaced with European naive AB serum to limit potential effect of human transmission blocking immunity and hence maximize the number of successfully infected mosquitoes (Gouagna et al., 2004). Serum was not changed in *An. coluzzii* experiments. As a negative control (uninfected mosquitoes), females were fed on the same blood in which gametocytes were heat-inactivated. Parasite inactivation was performed by placing the infectious blood in a thermo-mixer and heating at 43°C for 15 min and 900 rpm. This heat-inactivation prevents from infectiousness of gametocytes and does not affect the blood nutritive quality (Sangare et al., 2013). Mosquitoes exposed/unexposed to infection were therefore fed with blood from the same individual, avoiding the potential confounding effects of different blood origins on mosquito behaviours. Mosquito blood feeding was performed by distributing three hundred μl of blood in membrane feeders maintained at 37°C by water jackets; cups containing 60-80 mosquitoes were placed under the feeders to allow blood engorgement through Parafilm^®^ membranes for 2 hours. Fully blood-fed females were sorted out and placed in new cages (30x30x30 cm) where they had constant access to 5% glucose solution on cotton wool pads until the behavioural assays. A total of eight experimental replicates using 9 different gametocyte carriers were performed (see Table A1 for details).

### 2.3. Behavioural assays

A dual-choice olfactometer was used to study odour-mediated host choice by infected and uninfected mosquitoes (Lefèvre et al., 2009b; Lefèvre et al. 2010; Vantaux et al., 2015). *An. coluzzii* behavioural assays were carried out as in Vantaux et al. (2015). There was one slight modification in the set-up when used for *An. gambiae* and *An. arabiensis* assays for which the two tents were placed next to each other (Figure 1a). The olfactometer consisted of a source of two odours connected to two collecting boxes (30 × 30 × 40cm) linked by two glass tubes (L = 52cm, J = 10cm) to a downwind box (L × l × h=60 × 40 × 40cm) (Figure 1a). Odour stimuli (from human, calf or outdoor air) came from two tents connected to the two collecting boxes of the olfactometer by air vent hoses (Scanpart^®^, D × L = 10 × 300 cm). Gauze was placed at the junction between the air vent hose and the collecting boxes to prevent mosquitoes entering the tent. A fan was set up at the junction of the air vent hose and the tent to draw air from the tent to the collecting box and the downwind box. The wind speed in the two downwind arms was regulated at 15cm/s (±2cm/s) using a 435-4 Testo multi-functional meter (Testor, Forbach, France) equipped with a hot wire probe (range: 0 to 20m/s, accuracy: ± (0.03m/s +5% of mv)). The two tents were left outdoor while the collecting boxes and the downwind box were located inside an experimental room (Figure 1a).

**Figure 1.**
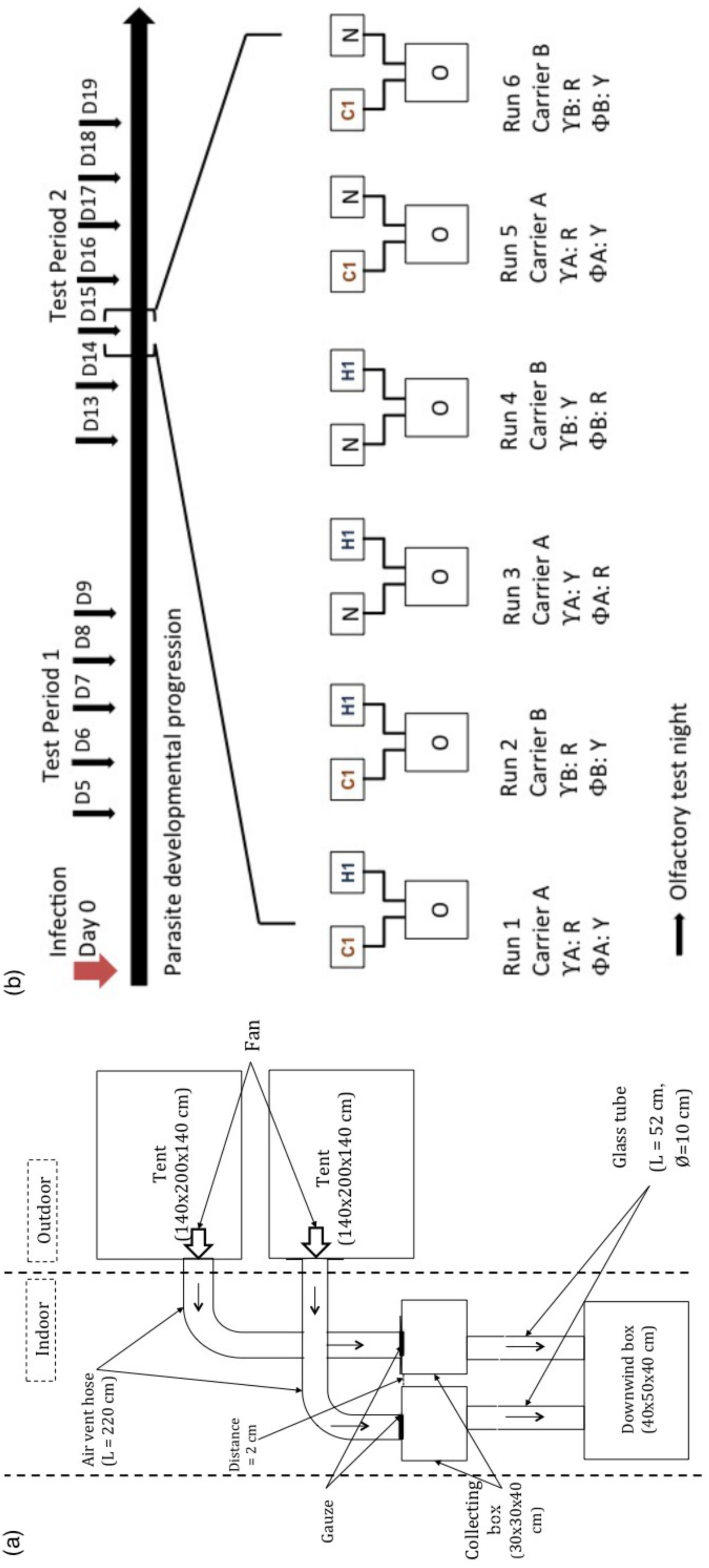
Schematic representations of (a) the dual-choice olfactometer and (b) the behavioural assays. C1 represents the tent containing calf 1, H1 represents the tent containing human volunteer 1, N represents the tent with outdoor air (control), O represents the Olfactometer, Y corresponds to mosquito infected status, Φ corresponds to mosquito uninfected status, R and Y represent the colours of the mosquitoes which are red and yellow respectively. The position of the tents was switched among replicates to account for side effect. Test period 1 and 2 correspond to the oocyst and sporozoite developmental stages in infected mosquitoes, respectively.

Mosquitoes were coloured with either red or yellow powders (Luminous Powder Kit, BioQuip) corresponding to their exposure status (received either an infectious blood-meal or a heat-inactivated uninfectious blood-meal) (Verhulst et al., 2013; Vantaux et al., 2015). The matching between exposure status and colours was switched between each run within a test day (Figure 1b). To increase mosquito response to host odours in the olfactometer set-up, the three mosquito species were deprived from glucose solution for 10 hours prior to the behavioural tests. During this period, mosquitoes had access to water only. For *An. gambiae* and *An. arabiensis* assays, in a single test day, a maximum of 6 runs each lasting 30 minutes were conducted using one mosquito species, and a total of 3 odour combinations: human *vs.* calf odour (H-C), human *vs.* outdoor air (H-O) and calf *vs.* outdoor air (C-O) (Figure 1b). For *An. coluzzii* experiments, only the human *vs.* calf odour (H-C) and human *vs.* outdoor air (H-O) combinations were carried out. In a single test day, 4 runs each lasting 30 minutes were carried out.

For each run, 20 uninfected controls and 20 gametocyte-exposed mosquitoes of similar age from the same experimental infection were simultaneously released in the downwind box of the apparatus. At the end of a run, the mosquitoes inside each of the two collecting boxes and the downwind box were removed with an aspirator, counted and kept in paper cups for subsequent analyzes (dissection of mosquito midgut and head/thorax, see below). Each batch of mosquitoes was tested once, so that a fresh batch of naive mosquitoes was used for each run. After each run, the olfactometer was washed with 70% alcohol to remove odour contaminants left from previous tests. Latex gloves were worn by the experimenter to avoid contamination of the equipment. The chronological order of the odour combinations was changed to eliminate possible confounding effect of odour combination and test time. The side of the tent connected to the collecting boxes was also switched to avoid positional effect. Different combinations of calves and humans were used as odour sources on each testing day to obviate any individual effect (a total of 21 volunteers and 18 calves). All volunteers who served as the human odour source were male Burkinabe around 20-30 years old who lived in Bobo Dioulasso. Calves of about similar size and weight as human volunteers were used to equalize quantity of emitted odours.

Mosquito host choice was tested at different time points corresponding to two distinct phases of the parasite development: (i) Test period 1 (5-8 days post-infection (dpi)) corresponding to the period of immature parasite development in the midgut (i.e. oocyst stage), (ii) Test period 2 (13-19 dpi) corresponding to the period of parasite transmission potential (when the sporozoites have invaded the mosquito salivary glands). The total number of run performed for each mosquito species, test period and odour combination is indicated in Table A1.

The day following behavioural testing, oocyst prevalence (proportion of *P. falciparum*-infected females) and intensity (number of oocysts in the midgut of infected females) were assessed by dissecting the midguts of mosquitoes that received an infectious blood-meal. Midguts were stained in a 1% mercurochrome solution and examined under a microscope (Vantaux et al., 2015). Heads and thoraces were used to determine sporozoite prevalence (proportion of infected females) by PCR assays for *An. coluzzii* females (Morassin et al., 2002) and by qPCR assays for the two other species (Boissière et al., 2013). Three groups of mosquitoes were thus obtained: (i) females that received a gametocyte-positive blood and became successfully infected; (ii) females that received a gametocyte-positive blood and remained uninfected; and (iii) females that received a heat-treated gametocytic blood (uninfected control).

### 2.4. Statistical analysis

We performed two sets of analyzes. In the first set, all data, including exposed-uninfected mosquitoes, were analyzed. In the second set, we excluded exposed-uninfected individuals to focus on the difference between infected and uninfected control mosquitoes. The two sets of analyzes yielded the same results, and, for the sake of clarity, only the second is reported in the main text (see Tables A2-A5 for the detailed output of the first set of analyzes).

All analyzes were performed in R (R Development Core Team, 2008). Binomial generalized linear mixed models (GLMMs) were fitted to investigate mosquito activation rate (proportion of mosquitoes caught in both collecting boxes out of the total number released) and odour choice (calculated separately for each odour combination, for example, human odour choice in the H-C combination was given as the proportion of mosquitoes entering human trap over the total mosquitoes entering both human and calf traps). In these models, infection treatment (two levels: infected and uninfected control), test period (two levels: test period 1 (oocyst stage) and test period 2 (sporozoite stage)), odour combination (three levels: H-C, H-O, and C-O for *An. gambiae* and *An. arabiensis,* and two levels: H-C, H-O for *An. coluzzii*) and relevant interactions were coded as fixed factors. Human volunteer and calf individual, and replicate were coded as random factors. Details of the fixed-effect and random-effect variables are shown in Tables A6 - A15. Because the *An. arabiensis* data were unbalanced (i.e. no data for carriers D and E during test period 1, see Table A1), activation rate and odour choice we analyzed separately for each test period. As there was complete separation of the data for one pair of human volunteer-calf individual in the H-C combination of test period 1 in the *An. arabiensis* data, volunteer was coded as a fixed factor in this host choice model only. We also verified for each combination, infection status and test period whether odour choice significantly differed from a random distribution between the two collecting boxes or whether mosquitoes displayed a statistically significant attraction to one odour.

For model selection, we used the stepwise removal of terms, followed by likelihood ratio tests (LRT). Term removals that significantly reduced explanatory power (*P <* 0.05) were retained in the minimal adequate model (Crawley, 2007). Post-hoc tests were carried out using the *testFactor* function in *phia* R package (Rosario-Martinez et al., 2015).

### 2.5. Ethical statement

Ethical approval was obtained from the Centre Muraz Institutional Ethics Committee (A003-2012/CE-CM) and National Ethics Committee of Burkina Faso (2014-0040). The protocol conforms to the declaration of Helsinki on ethical principles for medical research involving human subjects (version 2002) and informed written consent were obtained from all volunteers. This study was carried out in strict accordance with the recommendations in the Guide for the Care and Use of Laboratory Animals of the National Institutes of Health. The protocol was approved by both the Office of Laboratory Animal Welfare of US Public Health Service (Assurance Number: A5928-01) and national committee of Burkina Faso (IRB registration #00004738 and FWA 00007038). Animals were cared for by trained personnel and veterinarians.

## 3. Results

### 3.1. Infection

Of the 1448 *An. coluzzii* mosquitoes exposed to an infectious blood meal, 622 became infected (percentage ± 95% confidence interval: 42.96 ± 2.55 %, Figure A1), with a mean (± se) parasite intensity of 14.67 ± 1.41 (Figure A2). A total of 2161 *An. coluzzii* females (622 infected + 1539 uninfected control mosquitoes) were used for the analyzes. Of the 744 exposed *An. gambiae,* 570 were infected (76.61 ± 3.07 %, Figure A1), with a mean parasite intensity of 18.45 ± 1.71 (Figure A2). A total of 1260 *An. gambiae* females (570 infected + 670 control) were used for the analyzes. Finally, of the 447 exposed *An. arabiensis,* 243 were infected (54.36 ± 4.62%, Figure A1), with a mean parasite intensity of 8.77 ± 0.61 (Figure A2). A total of 694 *An. arabiensis* females (243 infected + 451 uninfected controls) were used for the analyzes. The differences in parasite prevalence and intensity observed across the different mosquito species can be explained by the use of different wild parasite isolates containing varying densities of gametocytes.

### 3.2. Activation rate

*Anopheles coluzzii.* Overall, 228 of 622 infected mosquitoes and 548 of 1539 control mosquitoes left the downwind box of the olfactometer to fly upwind into one of the two collection boxes (activation rate of 36.66 ± 3.79% and 35.61 ± 2.39%, respectively). *P. falciparum* infection did not affect *An. coluzzii* activation rate (χ^2^_1_ = 0.5, *P* = 0.48; Figure 2a). *An. coluzzii* displayed similar activation rate between the two test periods (χ^2^_1_ = 0.08, *P* = 0.78, Figure 2a) and the three odour combinations (χ^2^_1_= 3.21, *P* = 0.07, Figure 2a). Finally, there was no statistically significant interactions (test period x odour combination: χ^2^_1_= 1.06, *P* = 0.3; infection x odour combination: χ^2^_1_= 0.22, *P* = 0.64; infection x test period: χ^2^_1_= 113, *P* = 0.29; Table A6, Figure 2a).

**Figure 2.**
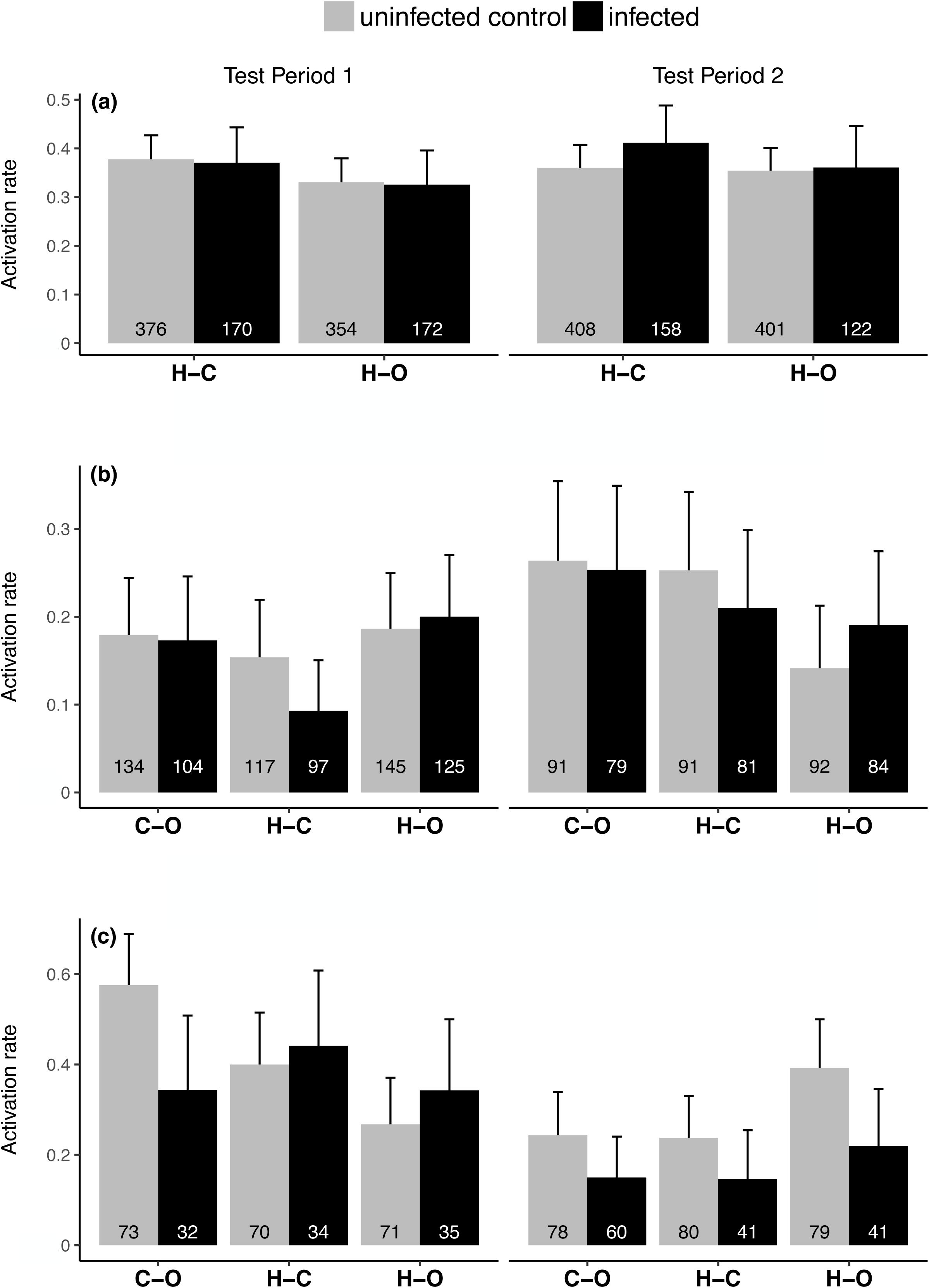
Mosquito activation rate, expressed as the proportion of mosquitoes caught in both collecting boxes out of the total number released in the downwind box for each treatment combination. (a) *Anopheles coluzzii,* (b) *Anopheles gambiae,* (c) *Anopheles arabiensis.* Numbers inside the bars indicate the total number of mosquitoes released across all runs. Error bars show the 95% confidence interval. C-O: for calf odour *vs* outdoor air combination, H-C: human odour *vs* calf odour combination, H-O: human odour *vs* outdoor air combination. Test Period 1 and Test Period 2 correspond to the oocyst and sporozoite stages in infected mosquitoes, respectively.

*Anopheles gambiae.* The overall activation rates of infected and uninfected control mosquitoes were 18.42 ± 3.18% (105/570) and 19.25 ± 2.99% (129/670) respectively.

Infection did not significantly affect *An. gambiae* activation rate (χ^2^_1_ = 0.22, *P* = 0.64, Figure 2b). There was a marginally significant effect of test period (χ^2^_1_ = 3.94, *P* = 0.047; Figure 2b). Although there was no significant main effect of odour combination on *An. gambiae* activation rate (χ^2^_2_ = 3.26, *P* = 0.2), there was a significant test period by odour combination interaction (χ^2^_2_= 8.44, *P* = 0.01, Table A7) such that the H-O odour combination induced the highest mosquito activation rate during test period 1 and the lowest during test period 2 (Figure 2b; Table A8). Finally, there was no infection by test period interaction (χ^2^_1_ = 0.23, *P* = 0.63), and no infection by combination interaction (χ^2^_2_ = 2.21, *P* = 0.33; Table A7).

*Anopheles arabiensis.* The activation rates of infected and uninfected control mosquitoes during test period 1 were 37.62 ± 9.45% (38/101) and 41.59 ± 6.6% (89/214) respectively. There was no influence of infection on *An. arabiensis* activation rate (χ^2^_1_ = 0.58, *P* = 0.45; Figure 2c; Table A9). There was a significant effect of odour combination on activation rate (χ^2^_2_ = 10.34, *P* = 0.005). In particular, mosquitoes were more activated in the C-O combination than in the H-O combination (post hoc tests: C-O vs. H-O: χ^2^_1_ = 9.96, *P* = 0.002; H-C vs. C-O: χ^2^_1_ = 1.79, *P* = 0. 18; H-O vs. H-C: χ^2^_1_ = 3.43, *P* = 0.06; Table A10). There was no significant interaction between infection and odour combination (χ^2^_2_= 4.95, *P* = 0.08; Table A9). The activation rate of infected *An. arabiensis* during test period 2 (16.90 ± 6.16%, n=142) was lower than that of uninfected individuals (29.11 ± 5.78%, n=237) (χ^2^_1_ = 6.69, *P* = 0.01; Figure c). We also found a significant effect of odour combination (χ^2^_2_ = 6.61, *P* = 0.04), such that mosquitoes were more activated in the H-O combination compared to the two other combinations (post hoc tests: H-O *vs.* H-C: χ^2^_1_ = 4.97, *P* = 0.03; H-O *vs.* C-O: χ^2^_1_ = 4.68, *P* = 0. 03; H-C *vs.* C-O: χ^2^_1_ = 0.01, *P* = 0.9; Table A11). We found no significant effect of infection by odour combination interaction (χ^2^_2_ = 0.18, *P* = 0.92 Figure 2c; Table A9).

### 3.3. Odour choice

*Anopheles coluzzii.* Infected and uninfected mosquitoes showed similar odour choice (H-C: χ^2^_1_ = 0.73, *P* = 0.4; H-O: χ^2^_1_ = 1.24, *P=* 0.43) with an overall attraction toward human odours of 71.70 ± 4.32% in the H-C and of 87.47 ± 3.42% in the H-O combinations (Figure 3a).

**Figure 3.**
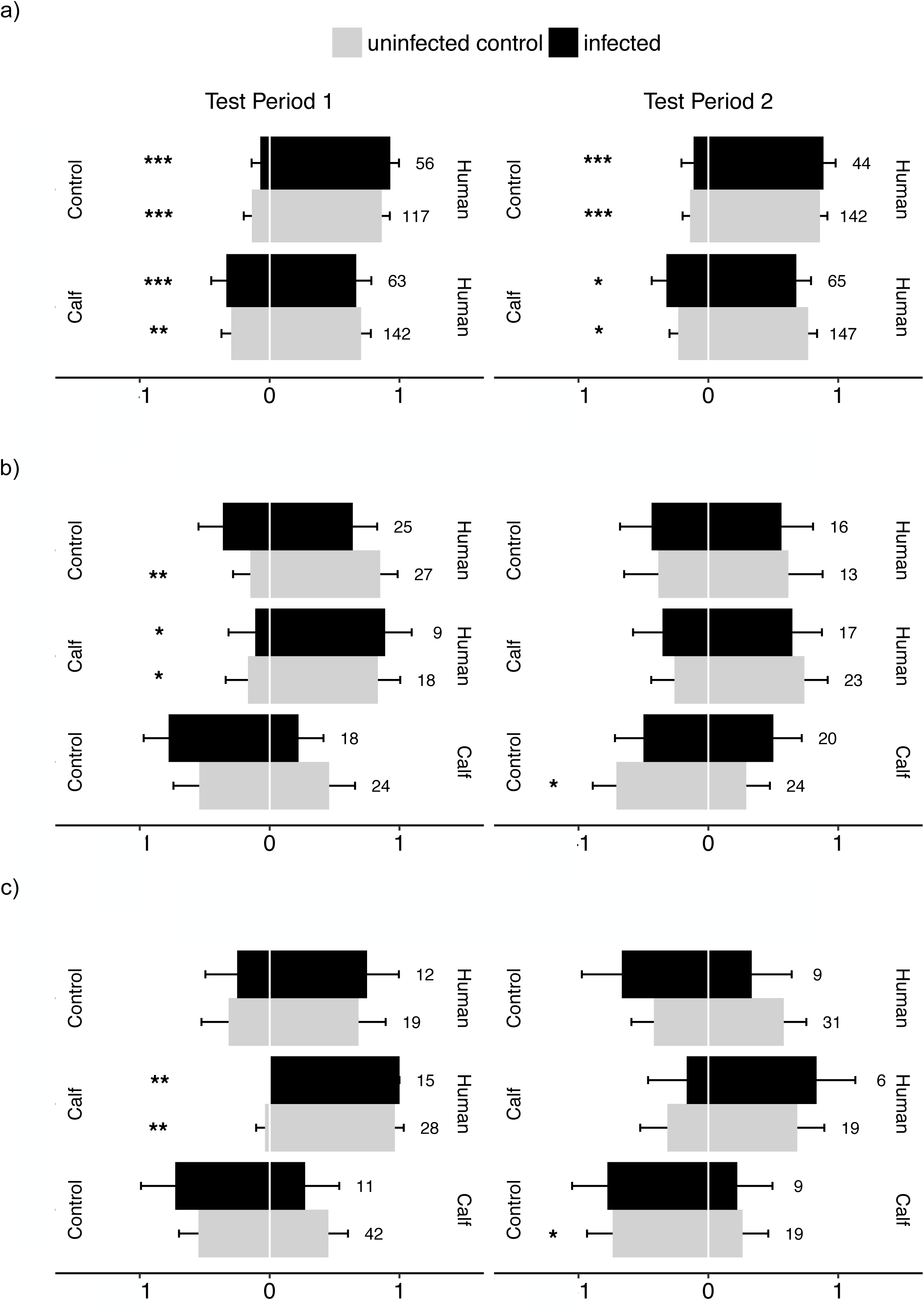
Mosquito odour-mediated choice - expressed as the proportion of mosquitoes caught in one collecting box out of the total number retrieved from both collecting boxes. (a) *An. coluzzii,* (b) *An. gambiae* (c) *An. arabiensis.* Data show proportion ± 95% confidence interval across all runs. Numbers indicate the total numbers of mosquitoes in both traps across all runs. The annotation human, calf, control corresponds to source of odour the mosquitoes chose Test Period 1 and Test Period 2 correspond to the oocyst and sporozoite stages in infected mosquitoes, respectively. (^*^ indicates significant bias toward an odour source; ^*^: *P* < 0.05, ^**^: *P* < 0.01, ^***^: *P* < 0.001).

There was no significant difference in odour choice between test period 1 and 2 in both the H-C (χ^2^_1_ = 0.03, *P* = 0.87) and the H-O odour combinations (χ^2^_1_ = 0.02, *P* = 0.89). Finally, there was no interaction between test period and infection in both the H-C (χ^2^_1_ = 0.31, *P* = 0.58) and H-O combinations (χ^2^_1_ = 0.62, *P* = 0.43; Figure 3a, Table A12).

*Anopheles gambiae.* In all three odour combinations, infected and uninfected mosquitoes displayed similar odour choice (H-C: χ^2^_1_ = 0.12, *P* = 0.73; H-O: χ^2^_1_ = 2.92, *P* = 0.09; C-O: χ^2^_1_ = 0.75, *P* = 0.75) with an overall attraction to human odours of 76.12 ± 10.21% in the H-C and 69.14 ± 10.06% in the H-O combinations, and a repulsion by calf odours of 62.79 ± 10.22% in the C-O combination (Figure 3b). There was no significant effect of test period (H-C combination: χ^2^_1_ = 1.96, *P* = 0.16; H-O combination: χ^2^_1_ = 1.66, *P* = 0.2; C-O combination: χ^2^_1_ = 0.4, *P* = 0.53) on *An. gambiae* odour choice. We found a significant infection by test period interaction for the C-O combination (χ^2^_3_ = 4.67, *P* = 0.03) with calf odours inducing a stronger repellence to infected mosquitoes than to uninfected mosquitoes during test period 1 and inversely during test period 2; Figure 3b, Table A13). Finally, there was no significant interaction between infection and test period in the H-C and H-O combinations (χ^2^_1_ = 0.47, *P* = 0.49 and χ^2^_1_ = 0.82, *P* = 0.37, respectively).

*Anopheles arabiensis.* For both test periods and all odour combinations, infected and uninfected mosquitoes displayed similar host choice (test period 1, H-C: χ^2^_1_= 0.87, *P* = 0.35; H-O: χ^2^_1_ = 0.16, *P* = 0.69; C-O: χ^2^_1_ = 1.2, *P* = 0.27; test period 2, H-C: χ^2^_1_ = 0.36, *P* = 0.55, HO: χ^2^_1_ = 1.39, *P* = 0.24, C-O: χ^2^_1_ = 0.06, *P* = 0.81; Figure 3c, Tables A14 and A15) with an attraction to human odours of 88.23 ± 7.66% in the H-C and 60.56 ± 11.37% in the H-O combinations and a repulsion by calf odour of 64.19 ± 10.44% in the C-O odour combination. There was no influence of volunteer/calf individual on odour choice (test period 1, H-C: χ^2^_1_ = 0.69, *P* = 0.41; Table A14).

## 4. Discussion

Several studies have demonstrated that malaria parasites can alter important behavioural features of their mosquito vectors in a stage dependant manner: oocyst-infected mosquitoes show reduced attraction to vertebrate odours and reduced avidity to feed, and, in contrast, sporozoite-infected individuals show enhanced attraction to vertebrate odours and enhanced avidity to feed (Hurd, 2003; Lefèvre and Thomas, 2008; Cator et al., 2012; Caljon et al., 2016). These behavioural alterations likely increase parasite transmission, provided that mosquito feeds are taken on a suitable vertebrate host species for the parasite. Our results indicate that, regardless of parasite developmental stage, *P. falciparum* infection did not alter mosquito activation rate, a surrogate of mosquito motivation to feed. While this finding contrasts with earlier studies (Rossignol et al., 1986; Cator et al., 2013; Smallegange et al., 2013), it supports two recent other studies, including one on *An. coluzzii* experimentally infected with sympatric wild isolates of *P. falciparum* (Cornet et al., 2013b; Vantaux et al., 2015), and suggests that manipulation of mosquito activity and response to host odours may not be a universal phenomenon. The expression of parasite-induced behavioural alterations, like any other phenotypic traits, may depend on local coevolutionary processes (Thompson, 2005). Hence, natural selection might not favour the evolution of manipulation in the studied populations if, for example, mosquito behaviour already ensures high parasite transmission or if the mosquito vector has evolved resistance (Daoust et al. 2015). Further investigations using sympatric and allopatric host-parasite combinations will be essential to integrate these local co-adaptation phenomena.

Our results also showed similar attractiveness of host odours to infected and uninfected mosquitoes. The overall expression pattern of mosquito host choice was consistent with a high degree of anthropophily in both infected and uninfected *An. coluzzii, An. gambiae* and *An. arabiensis.* The only significant effect of infection on mosquito odour choice was seen in sporozoite-infected *An. gambiae* which were less repelled by calf odours compared to oocyst-infected counterparts (Figure 3b). This result contrasts with the hypothesis of a parasite manipulation of mosquito odour preference and the expectation that sporozoite-infected individuals should be steered away from the odours of inappropriate vertebrate hosts. Although the precise reason behind this effect is currently unclear, it might be a mosquito response, and interactions among mosquito resources, infection and requirement of a blood-meal can be suspected. Because *Plasmodium* parasites utilize mosquito resources to develop and mature, sporozoite-infected *An. gambiae* females might be less choosy regarding the nature of the blood source, hence explaining this increased attraction to calf odour. At this stage, this remains speculative and further experiments are required to investigate the mechanisms underlying this effect and to explain why this pattern was not observed in *An. coluzzii* and *An. arabiensis.*

Our prediction was that sporozoites of *P. falciparum* may have evolved the ability to influence mosquito preference in a way that increases contact with humans - the appropriate vertebrate host of *P. falciparum* - by inducing in the mosquito vector a sensory bias for host odours that are correlated with suitability for the parasite. When simultaneously exposed to odours from calf and human, 71% and 76% of both sporozoite-infected and uninfected *An. coluzzii* and *An. gambiae* were retrieved from the human trap, hence confirming the anthropophilic behaviour of these species (Costantini et al. 1996; Costantini et al. 1999; Dekker et al. 2001;Takken and Verhulst 2013). Such anthropophily already ensures relatively high probability of transmission toward the appropriate host, perhaps making parasite manipulation of mosquito host choice useless in this system (i.e. weak selective pressures for its evolution). In other words, it is possible that, in this system, the balance between the transmission benefits of an increased preference for humans (e.g. from an anthropophily of 70 to 90%) and the costs associated with manipulation (Poulin, 1994) explain why this change has not evolved. Alternatively, it is possible that the parasite’s ability to modify its mosquito host choice did evolve but was not expressed here because of our experimental design.

First, while our olfactometer allows the study of long-range odour-mediated attractiveness, the full sequence of mosquito host-seeking process also includes short-range stimuli. Odour-mediated preference is critical at the initial step of host location; however final host decision might be influenced by cues other than odours, thereby determining alternative patterns of host preference. Indeed, olfactometer obviates stimuli such as visual cues, heat, moist, convective currents, and host movements. Under this scenario, sporozoite-infected *An. gambiae* would present similar host preference as uninfected counterparts in the early stages of the host-seeking process, when it mostly responds to host odours (i.e. long-range host preference), and then display increased attraction to humans at a shorter range, when other cues become more important (i.e. short-range host preference). In addition, we did not control the quantity of emitted host odour which could also affect mosquito behavioural responses (Costantini et al., 1996).

Second, our experiments were conducted between 6.30.pm and 11.30.pm while mosquito activation spans from 6.00.pm until early morning the following day with a peak of activity occurring around midnight. This activity peak is correlated with human resting behaviour to presumably maximize mosquito fitness (Lehane 2005). During this time frame, human are less defensive, possibly facilitating feeding and reduce mosquito mortality. Malaria parasites may have evolved the ability to finetune manipulation to the temporal behaviour of both the vectors and the vertebrate host. Under this scenario, manipulation of host choice might occur later in the night (e.g. during the peak of mosquito activity), and we hence may have missed it.

Third, *An. gambiae* and *An. coluzzii* mosquitoes from colonies continually replenished with F1 from wild-caught females were used here and it will also be important to use F0 from field mosquitoes since rearing insects in the laboratory for many generations is unlikely to represent the genetic diversity observed in nature.

Fourth, the uninfected control mosquitoes were fed on the same blood as infected mosquitoes but in which gametocytes were heat-inactivated. This procedure allows avoiding confounding effects of different blood origins on mosquito fitness and behavioural responses. However, these heat-killed gametocytes might trigger a mosquito immune response, which in turn, might affect mosquito behaviour. Although no study has, to our knowledge, directly explored whether heat-killed gametocytes can stimulate a mosquito immune response, there is evidence that alive gametocytes blood triggers a mosquito immune response that does not occur with heat-killed gametocytes (Mendes et al. 2008, 2011). In addition, the differences in immune gene expression between mosquitoes challenged with alive and dead *P. falciparum* gametocytes is similar to that observed between mosquitoes challenged with alive *P. berghei* gametocytes and mosquitoes that received a parasite-free blood-meal (Mendes et al. 2008, 2011). These results suggest that heat-killed *P. falciparum* gametocytes and parasite-free blood-meal may induce similar mosquito immune response. However, a study showed that a challenge with heat-killed *Escherichia coli* can generate mosquito behavioural changes similar to that observed in mosquitoes infected with rodent malaria parasites (Cator et al 2013). Overall, future studies should ideally include two control groups: heat-inactivated gametocytes from the same carrier (to avoid effects of different blood-meal sources on mosquito behaviour) and (ii) uninfected blood from a parasite-free donor (to avoid possible effects of heat-killed gametocytes on mosquito behaviour).

Finally, it is possible that the expression of parasite manipulation of host preference is more pronounced in some mosquito-parasite combinations than in others. In particular, we predicted that the expression of parasite manipulation of vector host choice might be more obvious in *An. arabiensis,* a presumably more zoophilic/opportunistic vector species (Takken and Verhulst, 2013). When simultaneously exposed to calf and human odours, we found that 90 *%* of *An. arabiensis* were retrieved from the human trap. This result contrasts with most existing studies on *An. arabiensis* host preference, which report an overall high degree of zoophily (Takken and Verhulst, 2013). Using odour-baited entry traps in Tanzania, Mahande et al. (2007), for example, showed a 90% zoophily in *An. arabiensis.* In contrast, of an estimated 1,800 field *An. arabiensis* collected using the same technique in Central Burkina Faso, slightly more than 8% were collected in the calf-baited trap, suggesting a high degree of anthropophily (Costantini et al., 1998). Our findings support the idea that West African populations of *An. arabiensis* may be generally more anthropophilic than East African ones (Costantini et al., 1999) and emphasizes that the anthropophilic/zoophilic label given to malaria mosquito species must be carefully interpreted and refer to populations rather than whole taxonomic unit.

We observed no parasite manipulation of mosquito odour-mediated host choice in the natural associations between *P. falciparum* and three of its major vector species, *An. coluzzii, An. gambiae,* and *An. arabiensis.* All three species were rather anthropophilic regardless of their infectious status. Further work is required to explore whether *P. falciparum* is able to modify its mosquito vertebrate choice in a way that increase transmission toward suitable host species. While our study examined the odour-mediated long-range mosquito host choice, determining the origin of blood-meals retrieved from uninfected, oocyst-infected and sporozoite-infected mosquitoes in the field may reveal the existence of specific manipulation. Future studies on specific manipulation in other vector systems would provide important information on the ecology and epidemiology of vector-borne diseases. A recent study suggested that the rodent-or bird-specialized *Borrelia* genospecies were unable to alter attraction of the generalist tick *Ixodes ricinus* to mouse odour (Berret and Voordouw, 2015). However, this study used ticks collected from the field and was not able to establish a causal relationship between *Borrelia* infection and attraction to mouse odour (Berret and Voordouw, 2015). Other possibly good model systems to study specific manipulation in vector-borne diseases are tsetse fly-transmitted trypanosomes. For example, *Glossina palpalis gambiensis* has a broad range of hosts in central Africa (humans, reptiles, bushbuck, and ox) and is the main vector of *Trypanosoma brucei gambiense* responsible for the medically important Human African trypanosomiasis. We would predict that once infected, flies are more attracted by human cues than by those of other vertebrates. Finally, parasite manipulation of mosquito host choice could theoretically occur at the intraspecific level (among different human individuals), with infected vectors biting more than expected less-immune hosts.

## Acknowledgments

This work was supported by the Agence Nationale de Recherche (grant number: 11-PDOC-006-01). We would like to thank all children and their parents for participating in this study, the local authorities for their support, as well as all the volunteers for the dual-port olfactometer assays. We are very grateful to the IRSS staff in Burkina Faso for technical assistance.

## References

Anderson, R.A., Koella, J.C., Hurd, H., 1999. The effect of *Plasmodium yoelii nigeriensis* infection on the feeding persistence of *Anopheles stephensi* Liston throughout the sporogonic cycle. Proc. Biol. Sci. 266: 1729–1733.

Berret, J., Voordouw, M.J., 2015. Lyme disease bacterium does not affect attraction to rodent odour in the tick vector. Parasite. Vectors. 8: 249.

Boissière, A., Gimonneau, G., Tchioffo, M.T., Abate, L., Bayibeki, A., Awono-Ambéné, P.H., Nsango, S.E., Morlais, I., 2013. Application of a qPCR assay in the investigation of susceptibility to malaria infection of the M and S molecular forms of *An. gambiae* s.s. in Cameroon. PLoS ONE. 8: e54820.

Caljon, G., De Muylder, G., Durnez, L., Jennes, W., Vanaerschot, M., Dujardin, J.C., 2016. Alice in microbes’ land: adaptations and counter-adaptations of vector-borne parasitic protozoa and their hosts. FEMS Microbiol. Rev. 40: 664–685.

Cator, L.J., George, J., Blanford, S., Murdock, C.C., Baker, T.C., Read, A.F., Thomas, M.B., 2013. ‘Manipulation’ without the parasite: altered feeding behaviour of mosquitoes is not dependent on infection with malaria parasites. Proc. Biol. Sci. 280: 20130711.

Cator, L.J., Lynch, P.A., Read, A.F., Thomas, M.B., 2012. Do malaria parasites manipulate mosquitoes? Trends Parasitol. 28: 467–470.

Cator, L., Lynch, P.A., Thomas, M.B., Read, A.F., 2014. Alterations in mosquito behaviour by malaria parasites: potential impact on force of infection. Malar. J. 13: 164.

Cornet, S., Nicot, A., Rivero, A., Gandon, S., 2013a. Malaria infection increases bird attractiveness to uninfected mosquitoes. Ecol. Lett. 16: 323–329.

Cornet, S., Nicot, A., Rivero, A., Gandon, S., 2013b. Both infected and uninfected mosquitoes are attracted toward malaria infected birds. Malar. J. 12: 179.

Costantini, C., Gibson, G., Sagnon, N., Della Torre, A., Brady, J., Coluzzi, M., 1996. Mosquito responses to carbon dioxide in a west African Sudan savanna village. Med. Vet. Entomol. 10: 220–227.

Costantini, C., Sagnon, N., della Torre, A., Coluzzi, M., 1999. Mosquito behavioural aspects of vector-human interactions in the *Anopheles gambiae* complex. Parassitologia. 41: 209–217.

Costantini, C., Sagnon, N., della Torre, A., Diallo, M., Brady J., Gibson G., Coluzzi, M., 1998. Odour-mediated host preferences of West African mosquitoes, with particular reference to malaria vectors. Am. J. Trop. Med. Hyg. 58: 56–63.

Crawley MJ. 2007. The R book. John Wiley & Sons Ltd, England.

Dabiré, R.K., Namountougou, M., Sawadogo, S.P., Yaro, L.B., Toé, H.K., Ouari, A., Gouagna, L.C., Simard, F., Chandre, F., Baldet, T., Bass, C., Diabaté, A., 2012. Population dynamics of *Anopheles gambiae* s.l. in Bobo-Dioulasso city: bionomics, infection rate and susceptibility to insecticides. Parasite. Vectors. 5: 127–135.

Dabiré, R.K., Sawadogo, P.S., Hien, D.F., Bimbilé-Somda, N.S., Soma, D.D., Millogo, A., Baldet, T., Gouagna, L.C., Simard, F., Lefèvre, T., Diabaté, A., Lees, R.S., Gilles, J.R., 2014. Occurrence of natural *Anopheles arabiensis* swarms in an urban area of Bobo-Dioulasso city, Burkina Faso, West Africa. Acta. Trop. 132: S35–41.

Daoust, S.P., King, K.C., Brodeur, J., Roitberg, B., Roche, B., Thomas, F., 2015. Making the best of a bad situation: host partial resistance and by-pass to behavioural manipulation by parasites? Trends Parasitol. 31: 413–418.

De Moraes, C.M., Stanczyk, N.M., Betz, H.S., Pulido, H., Sim, D.G., Read, A.F., Mescher, M.C., 2014. Malaria-induced changes in host odours enhance mosquito attraction. Proc Natl Acad. Sci. USA. 111: 11079–11084.

Dobson, A.P., 1988. The population biology of parasite-induced changes in host behaviour. Q. Rev. Biol. 63: 139–165.

Fanello, C., Santolamazza, F., della Torre, A., 2002. Simultaneous identification of species and molecular forms of the *Anopheles gambiae* complex by PCR-RFLP. Med. Vet. Entomol. 16: 461–464.

Gouagna, L.C., Bonnet, S., Gounoue, R., Verhave, J.P., Eling, W., Sauerwein, R., Boudin, C., 2004. Stage-specific effects of host plasma factors on the early sporogony of autologous Plasmodium falciparum isolates within *Anopheles gambiae*. Trop. Med. Int. Health. 9: 937–948.

Hien, D.F.S., Dabiré, K.R., Roche, B., Diabaté, A., Yerbanga, S., Cohuet, A., Yameogo, B.K., Gouagna, L.C., Hopkins, R.J., Ouedraogo, G.A., Simard, F., Ouedraogo, J.B., Ignell, R., Lefèvre, T., 2016. Plant-mediated effects on mosquito capacity to transmit human malaria. PLoS Pathog. 1–17.

Hughes, D.P., Brodeur, J., Thomas, F., 2012. Host Manipulation by Parasites (1st ed.). Oxford: Oxford University Press.

Hurd, H., 2003. Manipulation of medically important insect vectors by their parasites. Annu. Rev. Entomol. 48: 141–161.

Jacquin, L., Mori, Q., Pause, M., Steffen, M., Médoc, V., 2014. Non-specific manipulation of gammarid behaviour by *P. minutus* parasite enhances their predation by definitive bird hosts. PLoS ONE. 9: 1–12.

Kaldonski, N., Perrot-Minnot, M.J., Motreuil, S., Cézilly, F., 2008. Infection with acanthocephalans increases the vulnerability of *Gammarus pulex* (Crustacea, Amphipoda) to non-host invertebrate predators. Parasitology. 135: 627–632.

Koella, J.C., Packer, M.J., 1996. Malaria parasites enhance blood-feeding of their naturally infected vector *Anopheles punctulatus*. Parasitology. 113: 105–109.

Koella, J.C., Rieu, L., Paul, R.E.L. 2002. Stage-specific manipulation of a mosquito’s host-seeking behaviour by the malaria parasite *Plasmodium gallinaceum*. Behav. Ecol. 13: 816–820.

Koella, J.C., Sørensen, F.L., Anderson, R.A., 1998. The malaria parasite, *Plasmodium falciparum,* increases the frequency of multiple feeding of its mosquito vector, *Anopheles gambiae*. Proc. Biol. Sci. 265: 763–768.

Lacroix, R., Mukabana, W.R., Gouagna, L.C., Koella, J.C., 2005. Malaria infection increases attractiveness of humans to mosquitoes. PLoS Biol. 3: 1590–1593.

Lafferty, K.D., Morris, A.K., 1996. Altered behvavior of parasitized killifish increases susceptibility to predation by bird final hosts. Ecology. 77: 1390–1397.

Lagrue, C., Güvenatam, A., Bollache, L., 2013. Manipulative parasites may not alter intermediate host distribution but still enhance their transmission: field evidence for increased vulnerability to definitive hosts and non-host predator avoidance. Parasitology. 140: 258–265.

Lagrue, C., Kaldonski, N., Perrot-Minnot, M.J., Motreuil, S., Bollache, L., 2007. Modification of hosts’ behaviour by a parasite: Field evidence for adaptive manipulation. Ecology. 88: 2839–2847.

Lefèvre, T., Adamo, S.A., Biron, D.G., Missé, D., Hughes, D., Thomas, F., 2009a. Invasion of the body snatchers: the diversity and evolution of manipulative strategies in host-parasite interactions. Adv. Parasitol. 68: 45–83.

Lefèvre, T., Gouagna, L.C., Dabiré, K.R., Elguero, E., Fontenille, D., Costantini, C., Thomas F. 2009b. Evolutionary lability of odour-mediated host preference by the malaria vector *Anopheles gambiae*. Trop. Med. Int. Health. 14: 228–236.

Lefèvre, T., Gouagna, L.C., Dabiré, K.R., Elguero, E., Fontenille, D., Renaud, F., Costantini, C., Thomas, F., 2010. Beer consumption increases human attractiveness to malaria mosquitoes. PLoS ONE. 5: e9546.

Lefèvre, T., Koella, J.C., Renaud, F., Hurd, H., Biron, D.G., Thomas, F., 2006. New prospects for research on manipulation of insect vectors by pathogens. PLoS Pathog. 2: e72.

Lefèvre, T., Thomas, F., 2008. Behind the scene, something else is pulling the strings: Emphasizing parasitic manipulation in vector-borne diseases. Infect. Genet. Evol. 8: 504–519.

Lehane, M.J., 2005. The Biology of Blood-Sucking in Insects, Second Edition. New York, Cambridge University Press.

Lyimo, I.N., Ferguson, H.M., 2009. Ecological and evolutionary determinants of host species choice in mosquito vectors. Trends Parasitol. 25: 189–196.

Mahande, A., Mosha, F., Mahande, J., Kweka, E., 2007. Feeding and resting behaviour of malaria vector, *Anopheles arabiensis* with reference to zooprophylaxis. Malar. J. 6: 100.

Médoc, V., Beisel, J., 2009. Field evidence for non-host predator avoidance in a manipulated amphipod. Naturwissenschaften. 96: 513–523.

Médoc, V., Rigaud, T., Bollache L., Beisel, J., 2009. A manipulative parasite increasing an antipredator response decreases its vulnerability to a nonhost predator. Anim. Behav. 77: 1235–1241.

Mendes, A.M., Awono-Ambene, P.H., Nsango, S.E., Cohuet, A., Fontenille, D., Kafatos, F.C., Christophides, G.K., Morlais, I., Vlachou, D., 2011. Infection Intensity-Dependent Responses of *Anopheles gambiae* to the African Malaria Parasite *Plasmodium falciparum*. Infect. Immun. 79: 4708–4715.

Mendes, A.M., Schlegelmilch, T., Cohuet, A., Awono-Ambene, P., De Iorio, M., Fontenille, D., Morlais, I., Christophides, G.K., Kafatos, F.C., Vlachou, D., 2008. Conserved Mosquito/Parasite Interactions Affect Development of *Plasmodium falciparum* in Africa. PLoS Pathog. 4: e1000069.

Moore, J., 2002. Parasites and the behaviour of animals. New York: Oxford University Press.

Morassin, B., Fabre, R., Berry, A., Magnaval, J.F., 2002. One year’s experience with the polymerase chain reaction as a routine method for the diagnosis of imported malaria. Am. J. Trop. Med. Hyg. 66: 503–508.

Mouritsen, K.N., Poulin, R. 2003. Parasite-induced trophic facilitation exploited by a nonhost predator: A manipulator’s nightmare. Int. J. Parasitol. 33: 1043–1050.

Ngoubangoye, B., Boundenga L., Arnathau, C., Mombo, I.M., Durand, P., Tsoumbou, T., Otoro, B.V., Sana, R., Okouga, A.P., Moukodoum, N., Willaume, E., Herbert, A., Fouchet, D., Rougeron, V., Ba, C.T., Ollomo, B., Paupy, C., Leroy, E.M., Renaud, F., Pontier, D., Prugnolle, F., 2016. The host specificity of ape malaria parasites can be broken in confined environments. Int. J. Parasitol. 46: 737–744.

Ouédraogo, A.L., Guelbéogo, W.M., Cohuet, A., Morlais, I., King, J.G., Gonçalves, B.P., Bastiaens, G.J.H., Vaanhold, M., Sattabongkot, J., Wu, Y., Coulibaly, M., Ibrahima, B., Jones, S., Morin, M., Drakeley, C., Dinglasan, R.R., Bousema, T., 2013. A protocol for membrane feeding assays to determine the infectiousness of *P. falciparum* naturally infected individuals to *Anopheles gambiae*. Malar. World J. 4: 17–20.

Perkins, S.L., 2014. Malaria’s many mates: past, present, and future of the systematics of the order Haemosporida. J. Parasitol. 100: 11–25.

Poulin, R., 1994. The evolution of parasite manipulation of host behaviour: a theoretical analysis. Parasitology. 109: S109–S118.

Poulin, R., 2010. Parasite Manipulation of Host Behaviour. In edited by H. J. Brockmann, T. J. Roper, M. Naguib, K. E. Wynne-Edwards, J. C. Mitani, & L. W. Simmons (Eds.), Advances in the study of behaviour (pp.151–86). USA: Elsevier Inc.

Prugnolle, F., Durand, P., Ollomo, B., Duval, L., Ariey, F., Arnathau, C., Gonzalez, J.P., Leroy, E., Renaud, F., 2011. A fresh look at the origin of *Plasmodium falciparum,* the most malignant malaria agent. PLoS Pathog. 7: e1001283.

Rayner, J.C., Liu, W., Peeters, M., Sharp, P.M., Hahn, B.H., 2011. A plethora of *Plasmodium* species in wild apes: A source of human infection? Trends Parasitol. 27: 222–229.

R Development Core Team. 2008. R: A Language and Environment for Statistical Computing.

Rosario-Martinez, H. De, Fox, J., Team, R.C., 2015. Phia: Post-Hoc Interaction Analysis.

Rossignol, P.A., Ribeiro, J.M.C., Spielman, A., 1984. Increased intradermal probing time in sporozoite-infected mosquitoes. Am. J. Trop. Med. Hyg. 33: 17–20.

Rossignol, P.A., Ribeiro, J.M., Spielman, A., 1986. Increased biting rate and reduced fertility in sporozoite-infected mosquitoes. Am. J. Trop. Med. Hyg. 35: 277–279.

Roux, O., Vantaux, A., Roche, B., Yameogo, K.B., Dabiré, K.R., Diabaté, A., Simard, F., Lefèvre, T., 2015. Evidence for carry-over effects of predator exposure on pathogen transmission potential. Proc. Biol. Sci. 282: 20152430.

Sangare, I., Michalakis, Y., Yameogo, B., Dabire, R., Morlais, I., Cohuet, A., 2013. Studying fitness cost of *Plasmodium falciparum* infection in malaria vectors: validation of an appropriate negative control. Malar. J. 12: 2.

Santolamazza, F., Mancini, E., Simard, F., Qi, Y., Tu, Z., della Torre, A., 2008. Insertion polymorphisms of SINE200 retrotransposons within speciation islands of *Anopheles gambiae* molecular forms. Malar. J. 7: 163.

Schwartz, A., Koella, J.C., 2001. Trade-offs, conflicts of interest and manipulation in *Plasmodium-mosquito* interactions. Trends Parasitol. 17: 189–194.

Seppälä, O., Jokela, J., 2008. Host manipulation as a parasite transmission strategy when manipulation is exploited by non-host predators. Proc. Biol. Sci. 4: 663–666.

Seppälä, O., Karvonen, A., Valtonen, E.T., 2012. Behavioural mechanisms underlying ‘specific’ host manipulation by a trophically transmitted parasite. Evol. Ecol. Res. 14: 73–81.

Seppälä, O., Valtonen, E.T., Benesh, D.P., 2008. Host manipulation by parasites in the world of dead-end predators: adaptation to enhance transmission? Proc. Biol. Sci. 275: 1611–1615.

Smallegange, R.C., van Gemert, G.J., van de Vegte-Bolme, r M., Gezan, S., Takken, W., Sauerwein, R.W., Logan, J.G., 2013. Malaria Infected Mosquitoes Express Enhanced Attraction to Human Odour. PLoS ONE. 8: 8–10.

Takken, W., Verhulst, N.O., 2013. Host Preferences of Blood-Feeding Mosquitoes. Annu. Rev. Entomol. 433–453.

Thompson, J.N., 2005. The geographic mosaic theory of coevolution. USA: University of Chicago Press.

Vantaux, A., Dabiré, K., Cohuet, A., Lefèvre, T., 2014. A heavy legacy: offspring of malaria-infected mosquitoes show reduced disease resistance. Malar. J. 13: 442.

Vantaux, A., de Sales Hien, D.F., Yameogo, B., Dabiré, K.R., Thomas, F., Cohuet, A., Lefèvre, T., 2015. Host-seeking behaviours of mosquitoes experimentally infected with sympatric field isolates of the human malaria parasite *Plasmodium falciparum:* no evidence for host manipulation. Frontiers. Ecol. Evol. 3: 1–12.

Verhulst, N.O., Loonen, J.A., Takken, W., 2013. Advances in methods for colour marking of mosquitoes. Parasite. Vectors. 6: 200.

Wekesa, J.W., Copeland, R.S., Mwangi R.W., 1992. Effect of *Plasmodium falciparum* on blood feeding behaviour of naturally infected *Anopheles* mosquitoes in western Kenya. Am. J. Trop. Med. Hyg. 47: 484–488.

